# Obstructive Sleep Apnea Syndrome Disrupts Glymphatic-Related Physiological Brain Pulsations

**DOI:** 10.64898/2026.06.02.729357

**Authors:** Usko Huuskonen, Nuutti Santaniemi, Olli Juntunen, Teemu Myllylä, Vesa Korhonen, Matti Kinnunen, Tuomo Starck, Mirja Tenhunen, Sari-Leena Himanen, Mika Kallio, Janne Kananen

**Author notes:** These authors contributed equally to this work (Usko Huuskonen, Nuutti Santaniemi).

## Abstract

**Background:** Obstructive sleep apnea (OSA) affects over one billion people and increases neurodegenerative risk. Brain vasomotor, respiratory, and cardiac pulsations are thought to drive glymphatic clearance during sleep, yet OSA’s effect on these pulsations remains poorly understood.

**Methods:** We studied 20 healthy controls (HC; 39.7±8.0 y) and 12 patients with OSA (PWOSA; 53.0±11.0 y) using a four wavelength (690-980 nm) functional near-infrared spectroscopy (fNIRS) measuring oxygenated, deoxygenated and total hemoglobin, water and cerebrospinal fluid (HbO, HbR, HbT, H_2_O, CSF). Polar devices provided heart rate variability (HRV). Spectral power, coherence and phase transfer entropy analysis was performed for very low frequency (VLF), respiratory, and cardiac bands from 30 min sleep segments and event-based whole-night analysis assessed autonomic responses.

**Results:** OSA patients exhibited lower fNIRS spectral entropy for HbO, HbT, H_2_O & CSF (p<0.05), with increased VLF and respiratory band power across all concentrations (HbO, HbR, HbT, H_2_O & CSF, p<0.05). Signal coherence was reduced in respiratory and cardiac bands. Phase transfer entropy revealed disrupted directional coupling toward HbR (cardiac) and CSF-to-HbO (respiratory). HRV showed parallel VLF amplification (p <0.05), elevated heart rate during event-free and respiratory-event periods (p<0.001) and reduced Root Mean Square of Successive Differences (RMSSD, 32.5 vs 43.0 ms, p<0.001). Hypoxic burden correlated with cardiac band power of HbO, HbT and CSF (r>0.73).

**Conclusions:** OSA reorganizes cortical pulsatile dynamics – reducing complexity, amplifying low frequency power and suppfdsfsdfsdfsdisrupting directed coupling. This state may compromise sleep-related glymphatic clearance. fNIRS-based spectral analysis offers a promising bedside tool for monitoring brain pulsatility.

## 1 Introduction

Sleep is essential for maintaining full daily functionality, though if sleep is disrupted, the resulting effects can be adverse. Obstructive sleep apnea (OSA) is a globally prevalent cause of disturbed sleep, affecting at least one billion people worldwide (9–38%), depending on diagnostic criteria. Additionally, majority of the patients are unaware of their condition (Benjafield et al., 2019; Iannella et al., 2025; Senaratna et al., 2017). OSA is a disorder characterized by repeated obstruction of airflow during sleep, arising from complex mechanisms that narrow or collapse the upper airway (Canessa et al., 2011; White, 2005). This leads to intermittent oxygen deprivation, sleep fragmentation, and reduced sleep quality.

Recurrent episodes of hypoxia and sleep fragmentation associated with OSA initiate a cascade of pathological processes, including alterations in hemodynamics, heightened sympathetic activity, oxidative stress, systemic and neuroinflammation, and ultimately neuronal injury or loss (Lv et al., 2023). Patients with OSA exhibit pathological and functional changes in the brain, particularly in the hippocampus (Owen et al., 2019) and frontal lobe—regions vital for memory and cognition (Canessa et al., 2011; Owen et al., 2019). Furthermore, OSA has been associated with cognitive impairment and an increased risk of developing Alzheimer’s disease and Parkinson’s disease (Guay-Gagnon et al., 2022). Alzheimer’s patients with OSA have been found to have pathological biomarker levels (A*β*42 and t-tau) in the cerebrospinal fluid and accumulation of amyloid plaques even before dementia develops (Cui et al., 2022). The cumulative burden of nocturnal hypoxias may be captured more effectively by the hypoxic burden (HB) oxygen depletion load metric. The HB describes the total amount of decrease in oxygen saturation during sleep, taking into account the duration, depth and amount of hypoxia (Coso et al., 2024; Kulkas et al., 2013). The HB has been shown to predict heart failure with better accuracy than the traditional apnea-hypopnea index (AHI) (Azarbarzin et al., 2020).

The glymphatic system is a brain-wide clearance network, responsible for removing amyloid-β, tau-proteins and other neurotoxic metabolic byproducts from the brain (Bohr et al., 2022; Iliff et al., 2012). Current evidence suggests that this cleansing mechanism is modulated by three distinct physiological pulsations: cardiac, respiratory and very slow frequency (VLF), i.e. vasomotor waves (Helakari et al., 2022; Kiviniemi et al., 2016). The activity of the glymphatic system enhances during sleep, especially in the slow wave sleep (Hauglund et al., 2025; Hauglund and Nedergaard, 2025; Xie et al., 2013). During these stages, the diluted hydrodynamic expansion of the intercellular space within brain tissue is thought to reduce resistance to fluid flow, thereby facilitating the influx of cerebrospinal fluid (CSF) and promoting the efficient washout of metabolic waste (Hauglund et al., 2025; Hauglund and Nedergaard, 2025; Xie et al., 2013). Given the importance of these sleep dependent processes, disruptions in sleep architecture – such as those in OSA – may significantly compromise the brains metabolic maintenance.

In OSA, both apneic events and the reduction of slow-wave sleep may contribute to impaired glymphatic function (Roy et al., 2022). Thus, altered brain clearance may play an important role in the brain pathophysiology connected with OSA. Additionally, OSA disrupts autonomic nervous system (ANS) function, causing cardiovascular and physiological changes that persist beyond apneic events (Sequeira et al., 2019). Repetitive cycles of hypoxia, hypercapnia, and arousal trigger autonomic imbalance characterized by increased sympathetic activity and reduced parasympathetic tone (Sequeira et al., 2019). One way to assess sympathovagal balance is through heart rate variability (HRV), which is known to be significantly altered in patients with OSA (Qin et al., 2021).

Novel non-invasive techniques for studying the glymphatic system, such as functional near-infrared spectroscopy (fNIRS), makes it possible to study neurobiology in a new way (Gupta et al., 2026; Myllylä et al., 2018; Olopade et al., 2007; Yoon et al., 2025). Additionally, in previous NIRS examinations, abnormal deoxyhemoglobin levels were found, particularly during obstructive apnea events and impaired autoregulation of cerebral circulation (Olopade et al., 2007; Pizza et al., 2010). fNIRS can be used to perform an all-night high-time resolution monitoring of cerebral hemo- and hydrodynamics (Myllylä et al., 2018; Yoon et al., 2025). With this method, it is possible to accurately measure the physiological very low-frequency, respiratory, and cardiac pulsations that are thought to contribute glymphatic transport (Ilvesmäki et al., 2024; Kiviniemi et al., 2016; Väyrynen et al., 2026). Recent, preliminary fNIRS evidence shows that in healthy sleep, water signals and low-frequency oscillation power increase during N3 slow-wave sleep and correlate with EEG slow-wave activity, reflecting amplified vascular pulsations that support brain fluid exchange (Gupta et al., 2026).

Recent technological developments have enabled the high-fidelity monitoring of peripheral autonomic signals. For instance, optical sensors and photoplethysmography allow for continuous tracking of heart rate and oxygen saturation (Kinnunen et al., 2025), while mattress-type sensors can capture mechanical respiratory effort and ballistocardiographic dynamics (Tenhunen et al., 2013). Furthermore, impedance cardiography (ICG) provides non-invasive insights into cardiovascular dynamics and thoracic impedance fluctuations by measuring changes in electrical bioimpedance (Barbosa-Silva et al., 2005). Together, these non-invasive techniques offer a comprehensive window into cardiovascular and respiratory oscillations.

To our knowledge, there is little direct research data on how OSA affects physiological brain pulsations during sleep. Our hypothesis is that the pulsations are altered in OSA patients and reveal a possible connection to glymphatic impairment. In this explorative study setup, we wanted to make diverse spectral analysis of multimodal signals in time and frequency domain. Additionally, with wearable devices it would be possible to assess the effects of OSA and glymphatic dysfunction on the functioning of the autonomic nervous system.

## 2 Methods

### 2.1 Study Subjects

Patients with OSA and healthy control subjects were recruited to the multimodal sleep study (fNIRS, cardiorespiratory polygraphy, wearable devices), which was approved by the regional medical research ethics committee of the Wellbeing Services County of North Ostrobothnia (66/2023 & 21/2023) and Finnish medicine agency FIMEA (2023/003704). OSA patients were recruited from sleep apnea patients diagnosed at the Clinical Neurophysiology Department of Oulu University Hospital (Apnea-hypopnea index (AHI) > 5/h).

Control subjects were recruited via advertisement from university staff and students. All subjects were non-smokers aged 18–68 years without neurological diseases. Well-controlled comorbidities, such as hypertension managed with medication, were not exclusionary and were evaluated case-by-case. Subjects who were enrolled to the control group but turned to have sleep apnea referred to appropriate treatment, and were transferred to patient group in this study. The data were collected on hospital premises, with a member of the research team monitoring the measurements from an adjacent room. Written informed consent was obtained from all participants in accordance with the Declaration of Helsinki.

### 2.2 Measurement Protocol

Participants arrived at the sleep laboratory in the evening, approximately 1–1.5 hours before their usual bedtime. Upon arrival, participants completed all necessary consent forms and questionnaires. Laboratory assistants provided guidance for proper device placement. Once all devices were properly positioned and the initial signal quality verified, participants were allowed to engage in normal pre-sleep activities until ready to sleep. Acquisitions were started when participants indicated they were preparing to sleep, and recordings were synchronized across all systems to ensure temporal alignment.

Throughout the night, a study technician monitored recordings from an adjacent room with surveillance option to visually verify that measurement devices are on place and participant safety without disturbing the sleep. In the event of clear lead or sensor detachment, the study technician intervened to restore the signals with minimal patient contact.

### 2.3 Participants

Initially the study populations consisted of 44 individuals. After exclusions of five patients and seven control subjects due to poor signal quality, movement artifacts, technical failures, or insufficient sleep data, the groups selected for further analysis comprised of 20 healthy control subjects (HC_fnirs_; age 39.7±8.0 years, BMI 26.2±2.6 kg/m^2^, 5 females) and 12 patients with OSA (PWOSA_fnirs_; age 53.0±11.0 years, BMI 29.8±5.6 kg/m^2^, 1 female). The corresponding numbers for participants with successful wearable measurements were 18 healthy subjects (HC_polar_; age 40.7 ±8.9 years, BMI 26.1±2.8 kg/m^2^, 5 females) and 11 patients (PWOSA_polar_; age 53.4±12.4 years, BMI 30.2±6.3kg/m^2^, 1 female) participants. The same Polar groups were used for the event based analysis. The final sample size for each analysis varied by modality specific technical measurement issues such as unforeseen artefacts. The sample group forming flow is described in **Figure 1**.

**Figure 1.**
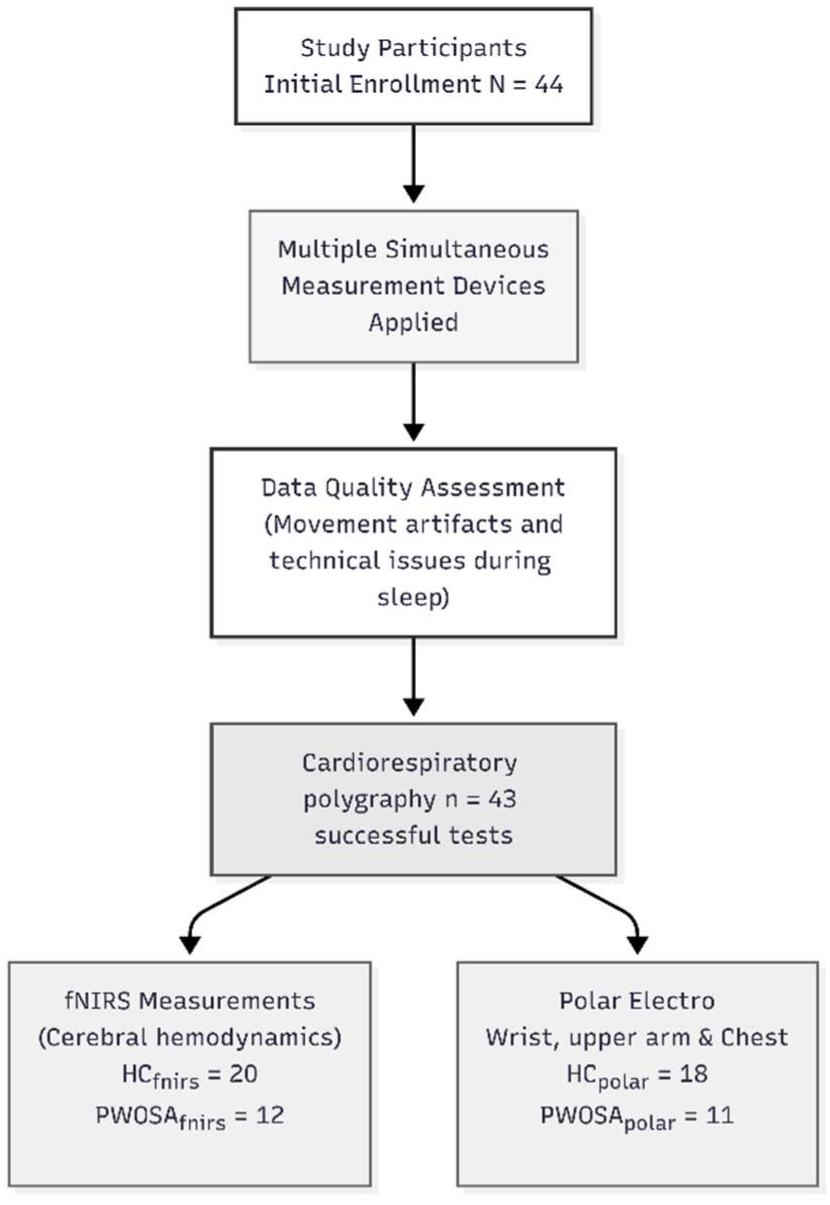
Flowchart of the study participants and group forming.

#### Cardiorespiratory polygraphy

The Nox T3s device (Nox Medical, ltd, Reykjavik) is a Type III cardiorespiratory polygraphy sleep apnea test adapted for laboratory use (Cairns et al., 2014). It records ventilatory effort through two respiratory inductive plethysmograph belts placed around the thorax and abdomen with sampling rate of 200Hz. Nasal airway pressure and flow are measured using nasal prongs with a sampling rate of 200 Hz. The device monitors body position and movement via a built-in 3D acceleration sensor. Audio and snoring are captured using a microphone with a 8kHz sampling rate, 3.5kHz bandwidth, 16-bit ADC. Oxygen saturation and pulse rate are measured using a Nonin WristOx2 Pulse Oximeter model 3150 BLE (sampling rate 75 Hz, storage rate 3 Hz) connected via Bluetooth. The registrations were analyzed by a specialist in clinical neurophysiology (UH), sleep assessed and respiratory events were scored according to AASM criteria (Malhotra, 2024). Hypoxic burden was derived from the SpO_2_ signal as the total are under the desaturation curve, effectively measuring the total load of low oxygen (Parekh, 2024).

### 2.4 Functional Near-Infrared Spectroscopy (fNIRS)

The study used an fNIRS device, designed at the University of Oulu and dedicated to sleep recordings. The methodology to calculated intracranial hydrodynamics is based on multi-wavelength near-infrared spectroscopy (NIRS) including four wavelengths (690, 810, 830, and 980 nm), where wavelength dependent attenuation follows the Beer–Lambert law. In particular, 980 nm is chosen due to its dominant absorbance of water, while 690, 810 and 830 nm are sensitive to oxyhemoglobin (HbO) and deoxyhemoglobin (Hb). From the raw NIRS signals calculated water (H2O) concentration contains both intracellular/extracellular water and blood-bound water. To isolate dynamical “free water” fluctuations associated mainly with CSF dynamics, the blood-related component was reduced based on total hemoglobin (HbT) calculations, as presented in (Myllylä et al. 2018). The remaining H2O (free) signal was interpreted as reflecting cerebrospinal fluid (CSF) dynamics in macro scale. The presented NIRS device has been used in numerous studies, recently e.g. to study cerebral water and hemodynamic response in whole brain radiotherapy (Miettinen et al., 2025) and brain aging (Ilvesmäki et al., 2025).

The battery-powered wearable device consists of a multiwavelength light source unit positioned at the center of the forehead and two photodetectors placed bilaterally on the forehead at a source–detector distance of 3 cm. Each emitted wavelength is individually modulated, enabling the photodetectors to distinguish the corresponding backscattered and reflected optical signals originating from brain tissue at a sampling rate of 250 Hz. To enhance sensitivity to intracranial hydrodynamics and reduce the influence of superficial scalp signals, additional measurements were acquired at a short source–detector separation of 1 cm. These superficial signals were subsequently subtracted from the 3 cm measurements to improve detection of hemodynamic and water-related dynamics originating from deeper regions.

### 2.5 Polar Devices

The following devices were employed from Polar Electro Oy (Kempele, Finland) to monitor ANS function, movement, and cardiovascular dynamics:

- **Polar Elixir Biosensing Solution:** A wrist-worn device (Vantage V3) utilizing optical measurement (photoplethysmography, PPG) at wavelengths of 550 nm (green), 660 nm (red), and 920 nm (infrared), combined with a 3D acceleration sensor. Sampling rate: 14 Hz for optical signals, 50 Hz for inertial measurements. This device provided validated SpO_2_ measurements (Russell et al., 2024) in addition to heart rate variability (HRV) metrics serving as indicators of autonomic nervous system function. This device was the primary source of Polar data in 30 min sleep segment analysis and event based whole-night analysis.
- **Multiwavelength Optical Sensor:** An upper arm wearable device performing optical measurements (PPG) at 550 nm, 660 nm, and 920 nm, integrated with a 3D acceleration sensor. Sampling rate: 14 Hz for optical signals, 50 Hz for inertial measurements. This device provided validated SpO_2_ measurement (Kinnunen et al., 2025). Heart rate data from this device were use in the event-based whole night HR analysis in Figure 9.
- **Impedance Cardiography Device:** A chest-worn strap measuring single-lead electrocardiography (ECG) and bioimpedance, equipped with a 3D acceleration sensor. Sampling frequency: 130 Hz for ECG and impedance, 50 Hz for inertial sensors. In this study, this device was used to Bioimpedance Phase Angle, serving as indicator of autonomic nervous system function and cellular integrity.

### 2.6 Data Analysis

Two complementary analysis approaches were employed: (1) synchronized 30-minute segment analysis combining fNIRS, cardiorespiratory polygraphy and Polar data from identical time windows, and (2) event-based whole-night analysis using Cardiorespiratory polygraph and Polar data to assess autonomic responses during respiratory events.

#### 2.6.1 30-Minute Sleep Segment Analysis

As the used multimodal setup and normal movements during sleep can introduce artifacts, we ensured high-quality data by having an expert validate the recordings and selecting, for each participant, a 30-minute segment in which both the fNIRS and cardiorespiratory polygraph signals were simultaneously of good quality. For each subject, a 30-minute segment was selected in which all fNIRS signals met high-quality standards with minimal artifacts. The corresponding Polar data from the same time window was extracted using synchronized timestamps.

##### fNIRS Spectral Analysis

To streamline further analysis, the concentration signals were downsampled to 50 Hz. A 0.005 Hz high-pass filter was utilized to remove slow DC drifts, ensuring signal stability. Signal complexity was then quantified via spectral entropy (Shannon, 1948) within a 0.005–1.5 Hz window. This range effectively encapsulates all relevant physiological rhythms, from very low-frequency vasomotion to cardiac pulses, without introducing higher frequency components that were not under interest in current study.

Additionally, for further frequency domain analysis, fNIRS data were further band passed for three physiological brain pulsations frequencies. Therefore, spectral power density was calculated for three frequency bands across all five concentrations (HbO, HbR, HbT, H_2_O, CSF) and for HRV:

- Very low frequency (VLF_fNIRS_): 0.01–0.1 Hz
- Respiratory: 0.1–0.5 Hz
- Cardiac: 0.5–1.5 Hz

With these frequency bands every concentration signals spectral power densities were calculated and compared statistically between groups (15 different comparisons).

##### Polar HRV Analysis

Heart rate variability was analysed from pulse-to-pulse intervals (ms, PPI) derived from the Polar Vantage V3 wrist device. Frequency domain analysis was performed to calculate power and entropy in the same frequency bands as fNIRS. PPI’s were resampled to 4 Hz using cubic spline interpolation prior to spectral analysis as per HRV Task Force recommendations (Task Force ESC/NASPE, 1996) and power spectral density (PSD) was estimated using Welch’s method (Hamming window, segment length 1024 samples). Normalized (H / log_2_(n)) Shannon entropy (Shannon, 1948) was used as a measure of signal regularity. The previously described frequency bands (VLF_fNIRS_, Respiratory and cardiac) were used for HRV calculations.

##### HRV Specific analysis

HRV metrics were also analysed in standard HRV frequency bands (Task Force ESC/NASPE, 1996):

- Very low frequency (VLF_HRV_): 0.003–0.04 Hz
- Low Frequency (LF): 0.04–0.15 Hz
- High Frequency (HF): 0.15–0.4 Hz
- LF/HF Ratio

Time domain analysis included the root mean square of successive differences (RMSSD) calculated using 10-sample sliding windows.

##### Connectivity Analysis

The following comparisons were selected for coherence and phase transfer entropy and the analysis was done within the same three frequency bands introduced earlier. In cardiac band the comparison pairs were selected to test for the cerebral windkessel (CWK) effect, the damping of arterial pulsations by the craniospinal environment (Wagshul et al., 2011; Westerhof et al., 2009) and to assess the arterial to venous coupling. All concentration signals HbO, HbT, CSF & H_2_O were compared against HbR. HbR was selected since it can be considered as proxy for venous compartment (Obrig and Villringer, 2003) and arterial pulsations are thought to give some pulsatile nature to venous outflow (Iliff et al., 2012; Kiviniemi et al., 2016; Oliveira et al., 2025; Wagshul et al., 2011). The CSF and H_2_O comparisons target the CWK mechanism, while HbO and HbT comparisons assess also the arterial to venous coupling. The HbT/HbO pair was included to assess arterial to venous coupling, examining whether total hemodynamic concentration carries some information that is beyond the oxygenated fractions. The CSF /HbO pair was selected for both cardiac and respiratory bands: in the cardiac band to capture CSF-driven venous outflow, and in the respiratory band to represent for CWK effect at lower-frequency pressure swings. In respiratory band the comparisons were selected to analyse for the thoracic pump on intracranial fluid displacement (Dreha-Kulaczewski et al., 2015). The HbT / CSF & HbT/H_2_O pairs were selected to account for volumetric balance and fluid movement. The HbT/HbO pair selected to account for the thoracic pump. In the VLF_fNIRS_ band we selected to compare HbT / CSF and HbT/H_2_O to be able to assess the slow wave fluid movement (Helakari et al., 2022). The HbT/HbR comparison was included to capture the neurovascular coupling (Kiviniemi et al., 2016). All the 13 comparisons are presented below:

- Cardiac band:

`o` CSF/HbR (venous-CSF coupling)
`o` HbO/HbR (arterial venous coupling)
`o` HbT/HbR (total venous pulsatility)
`o` H_2_O/HbR (parenchyma compliance)
`o` HbT/HbO (venous fraction)
`o` CSF/HbO (CSF driven venous)
- Respiratory band:

`o` HbT/CSF (thoracic pump)
`o` HbT/H_2_O (volumetric balance)
`o` HbT/HbO (respiratory oxygenation)
`o` CSF/HbO (respiratory windkessel)
- VLF_fNIRS_ band:

`o` HbT/CSF (slow wave fluid movement)
`o` HbT/H_2_O (slow wave fluid movement)
`o` HbT/HbR (neurovascular coupling)

##### Coherence analysis

In order to assess the degree of linear synchrony between fNIRS concentration signals, magnitude-squared coherence was calculated. Spectral densities were estimated using Welch’s method. This is consistent with the band power method described above. Coherence is a symmetric and undirected measure. It quantifies the strength of coupling between two signals but provides no information about the direction of that interaction. Therefore, directional coupling was assessed separately using phase transfer entropy for the same concentration signal pairs for 30min signals.

##### Phase Transfer Entropy (pTE)

Directed coupling between fNIRS concentration signals was assessed using phase transfer entropy (PTE). The computation was done as described by (Lobier et al., 2014). The instantaneous phase were extracted from each band-pass filtered signal with Hilbert transform. *PTE*_X→Y_ = *H*(*θ*_Y_, *θ*′_Y_) + *H*(*θ*′_Y_, *θ*′_X_) − *H*(*θ*^r^_Y_) − *H*(*θ*_Y_, *θ*′_Y_, *θ*′_X_). In the equation θ denotes instantaneous phase. Primes denote the past state at analysis lag (sigma). *H* denotes Shannon entropy. It was estimated by adaptive histogram binning based in the circular standard deviation of each phase series. A value of 0 indicates no directed coupling. The net directionality of coupling was quantified as Δ*TE* = *PTE*_X→Y_ − *PTE*_Y→X_ Positive values indicate net information flow form X to Y. Negative values indicate the reverse (Väyrynen et al., 2026). Analysis lags were selected to reflect the expected propagation delay within each frequency band. The lags were 500 sampled (10 s) for VLF_fNIRS_, 250 samples (5 s) for the respiratory band and 33 samples (0.66 s) for the cardiac band.

All fNIRS preprocessing was performed in MATLAB 2025b; Polar data analysis was performed in Python 3.13 (Python Software Foundation, 2026).

#### 2.6.2 Event-Based Whole-Night Analysis

Using the complete overnight Polar recordings (HC_polar_ & PWOSA_polar_), physiological responses on heart rate (HR), HRV, the depth of the valley of RRI fluctuation (VRRI) and Bioimpedance phase angle (PhA) were compared between OSA patients and control subjects during different respiratory events:

- Respiratory event periods (hypopnea & obstructive apnea events, desaturation events)
- Non-respiratory event periods (referred to as *Event-free*)

The VRRI was calculated as described by (Arikawa et al., 2020). This dual window approach distinguishes acute apnea-induced heart rate changes from slower baseline drifts.

##### Bioimpedance Analysis

PhA measured at 65 kHz, reflects the ratio of tissue reactance to resistance and is influenced by cellular membrane capacitance and tissue hydration (Holder, 2004). At this frequency, the injected current partially penetrates cell membranes, making PhA sensitive to changes in both extracellular and intracellular compartments (Holder, 2004). OSA is associated with chronic systemic inflammation and oxidative stress (Lv et al., 2023), which may alter tissue properties detectable by thoracic bioimpedance. We utilized the Polar chest strap to measure PhA during sleep as an exploratory metric to assess whether thoracic tissue electrical properties differ between OSA patients and control subjects during overnight recordings.

#### 2.6.3 Statistical Analysis

Group differences in participant characteristics (Table 1) were assessed with Welch’s two sample t-test. All subsequent between-group comparisons were conducted using non-parametric Mann-Whitney U tests, with significance set at p *<* 0.05. fNIRS statistics were computed in Origin 2025b; Polar statistics were computed in Python using SciPy and similar Mann-Whitney U test.

**Table 1.**
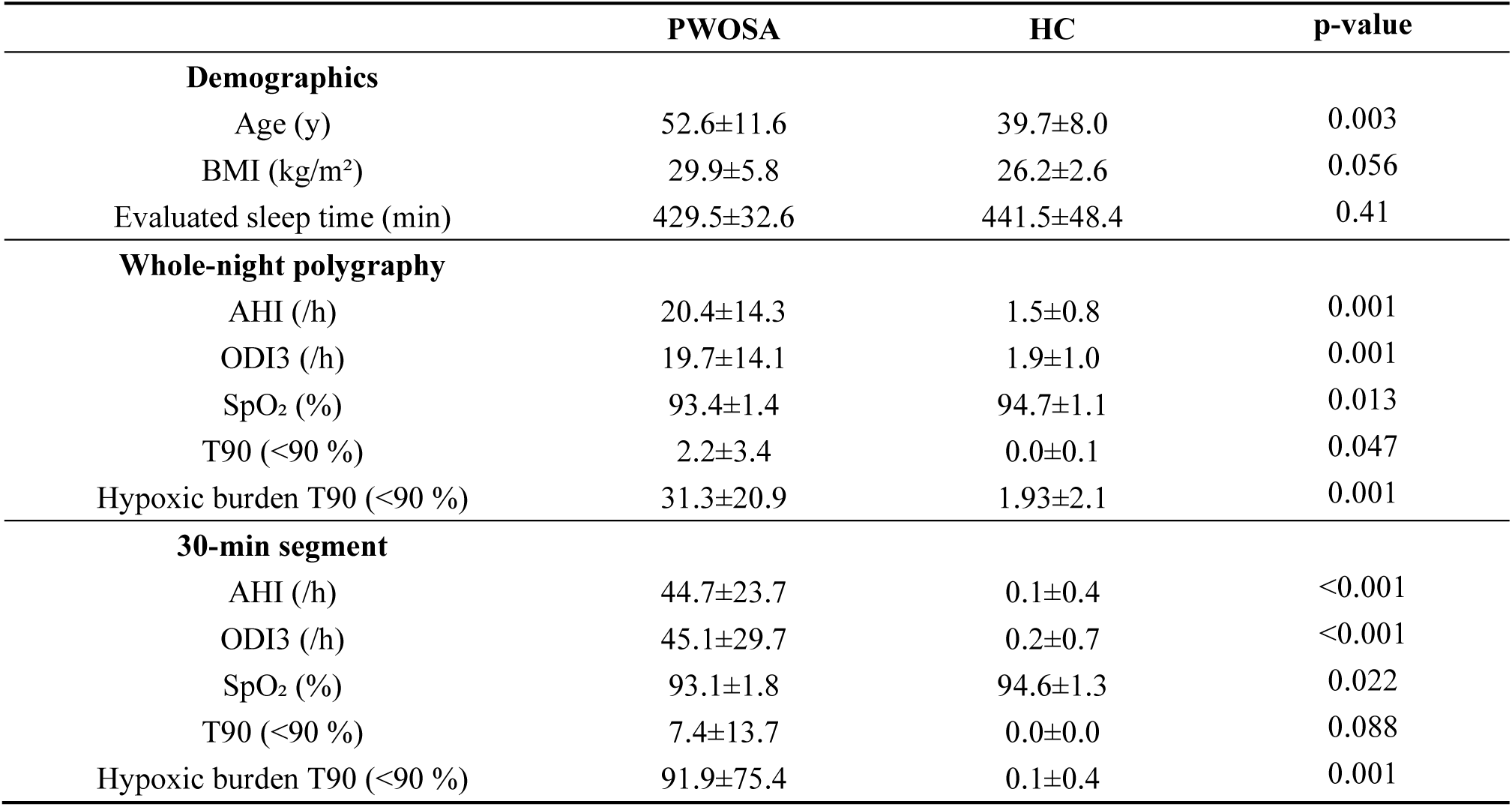
Comparison of participant characteristics and sleep related metrics between groups. Values represented as mean ± SD. Statistical comparisons were performed via Welch’s two sample t-test.

|  | PWOSA | HC | p-value |
| --- | --- | --- | --- |
| **Demographics** |  |  |  |
| Age (y) | 52.6±11.6 | 39.7±8.0 | 0.003 |
| BMI (kg/m<sup>2</sup>) | 29.9±5.8 | 26.2±2.6 | 0.056 |
| Evaluated sleep time (min) | 429.5±32.6 | 441.5±48.4 | 0.41 |
| **Whole-night polygraphy** |  |  |  |
| AHI (/h) | 20.4±14.3 | 1.5±0.8 | 0.001 |
| ODI3 (/h) | 19.7±14.1 | 1.9±1.0 | 0.001 |
| SpO<sub>2</sub> (%) | 93.4±1.4 | 94.7±1.1 | 0.013 |
| T90 (<90 %) | 2.2±3.4 | 0.0±0.1 | 0.047 |
| Hypoxic burden T90 (<90 %) | 31.3±20.9 | 1.93±2.1 | 0.001 |
| **30-min segment** |  |  |  |
| AHI (/h) | 44.7±23.7 | 0.1±0.4 | <0.001 |
| ODI3 (/h) | 45.1±29.7 | 0.2±0.7 | <0.001 |
| SpO<sub>2</sub> (%) | 93.1±1.8 | 94.6±1.3 | 0.022 |
| T90 (<90 %) | 7.4±13.7 | 0.0±0.0 | 0.088 |
| Hypoxic burden T90 (<90 %) | 91.9±75.4 | 0.1±0.4 | 0.001 |

## 3 Results

### 3.1 Participant Characteristics

The patients (PWOSA) were older and had higher BMI (Table 1) than the control subjects. Mean sleep duration did not differ significantly.

The control group’s whole-night apnea-hypopnea index (AHI) was 1.6±1.0, and the patients’ AHI was 18.6±14.0. For the 30-minute high-quality segments used in spectral analysis, equivalent AHI values were 0.1±0.4 in HC and 44.7±23.7 in PWOSA. Table 1 summarizes participant demographics and sleep polygraphy parameters.

### 3.2 30-Minute Segment Analysis

#### fNIRS Spectral Entropy

Spectral entropy analysis revealed a distinct loss of physiological complexity in sleep apnea. PWOSA_fnirs_ group showed significantly lower entropy in HbO, HbT, H_2_O & CSF signals relative to control subjects (0.0012 *<* p *<* 0.0033, Figure 2 A), whereas HbR complexity was somewhat preserved (p = 0.07; Figure 2A). Interestingly, this loss of complexity was echoed in the autonomic data. Although total-band HRV entropy (0.01–1.5 Hz) was unchanged, the entropy of the VLF_fNIRS_ component (0.01– 0.1 Hz) was significantly reduced in the OSA group (p *<* 0.05). This concordant decrease in VLF_fNIRS_ entropy across both cerebral and cardiac measurements suggests a systemic dampening of physiological variability Figure 3 B1.

**Figure 2.**
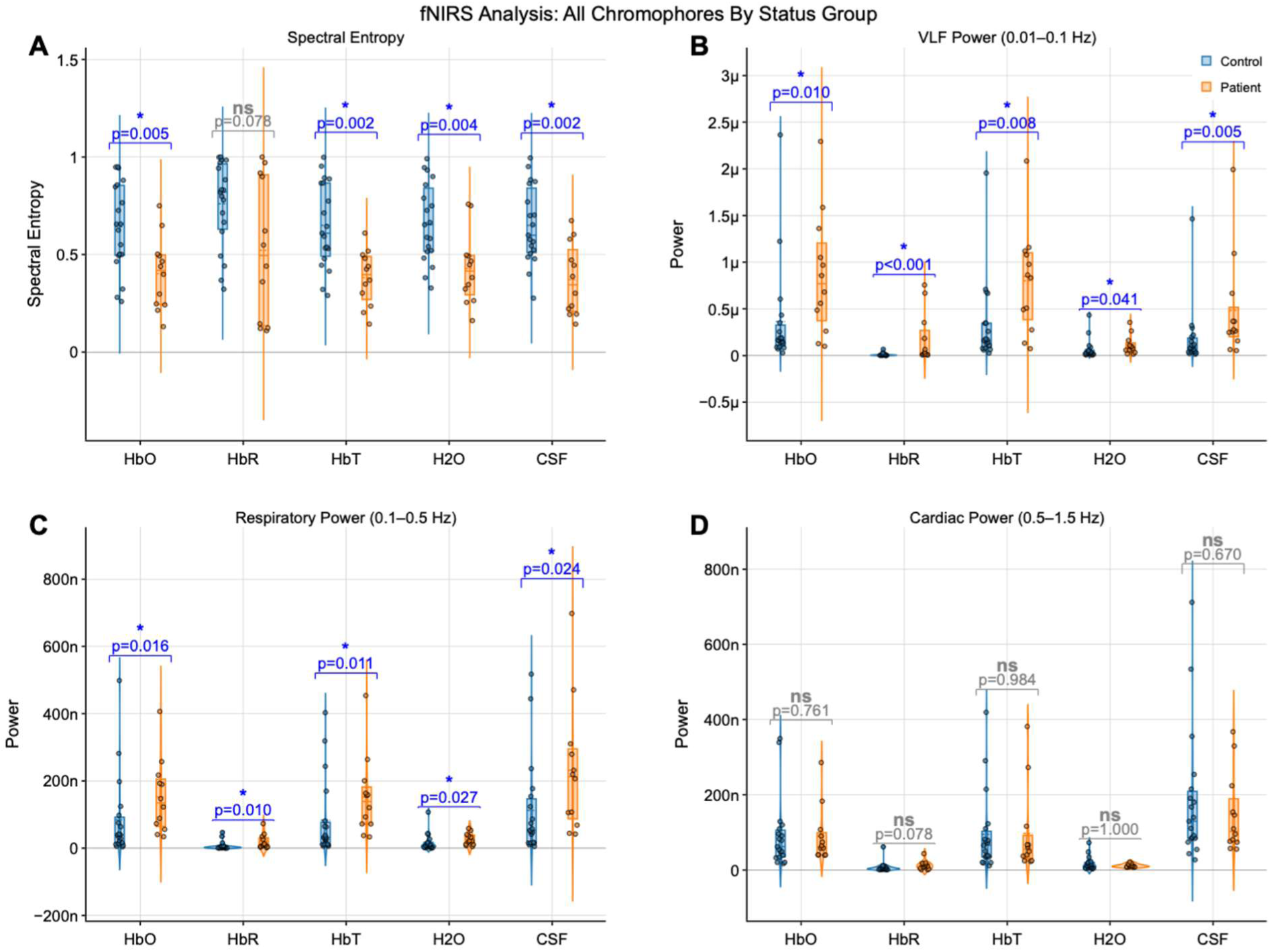
fNIRS spectral analysis during 30-minute high-quality segments in HC_fnirs_ and PWOSA_fnirs_ groups. (A) Spectral entropy was significantly lower in OSA patients for HbO, HbT, and H_2_O. (B) VLF band power (0.01–0.1 Hz) was significantly elevated in OSA patients across all concentrations. (C) Respiratory band power (0.1–0.5 Hz) showed significant increases in OSA patients. (D) Cardiac band power (0.5–1.5 Hz) did not differ between groups. *p < 0.05

**Figure 3.**
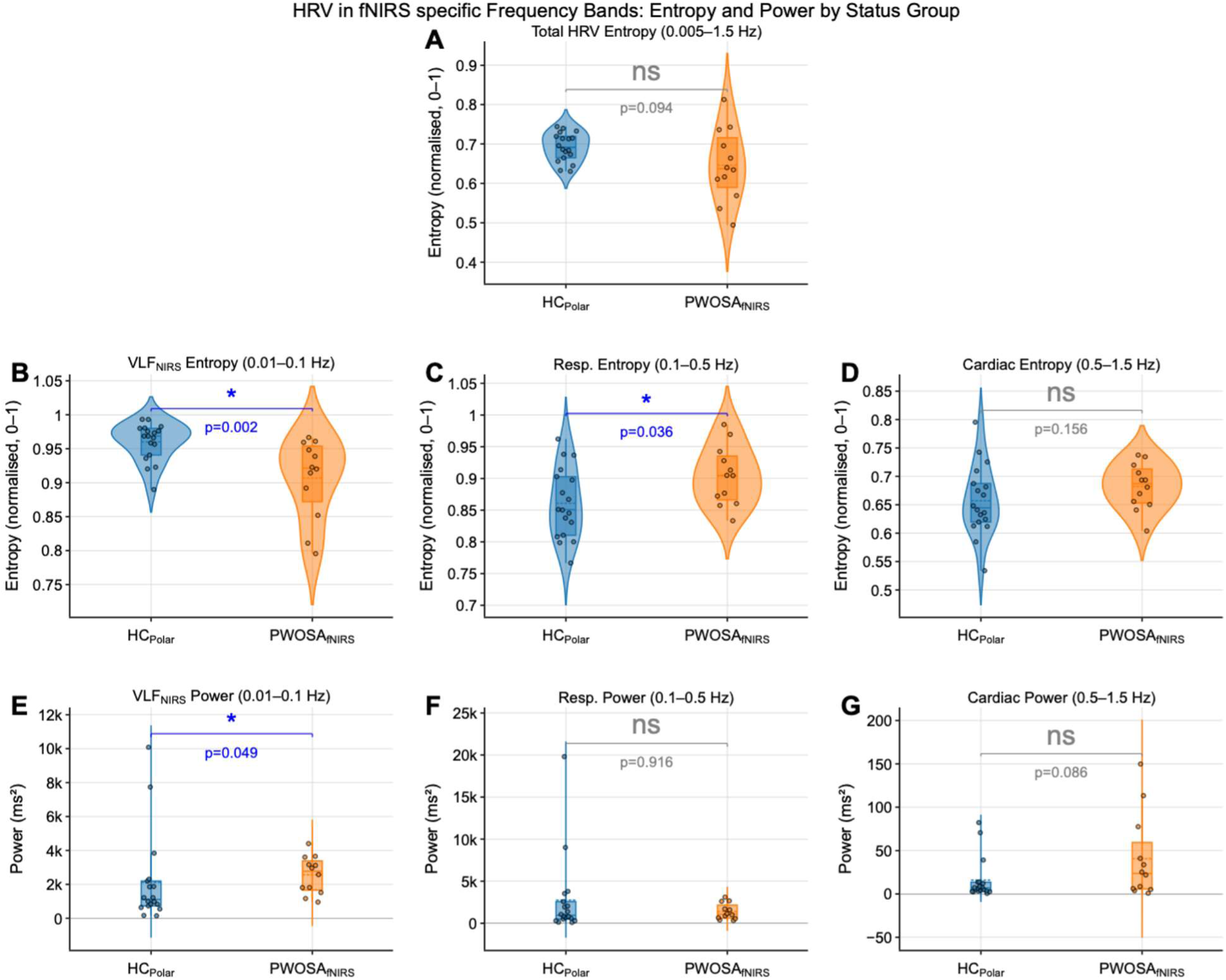
Spectral entropy of HRV in HC_polar_ and PWOSA_polar_ groups. The VLF_fNIRS_ band (0.01–0.1 Hz) showed significantly lower entropy in OSA patients.

#### Spectral Power Density

Spectral power density analysis of fNIRS and Polar HRV signals revealed a consistent increase in power within the VLF_fNIRS_ and respiratory bands across all four concentrations in the OSA group:

- **VLF_fNIRS_ band (0.01–0.1 Hz):** All concentrations (HbO, HbR, HbT, H_2_O, CSF) were significantly higher in the patient group with p values below p *<* 0.05 (Figure 2B). The similar phenomenon was observed with HRV power density Figure 3 B2.
- **Respiratory band (0.1–0.5 Hz):** All concentrations were significantly higher in the patient group. For H_2_O, the respiratory-band increase was significant at p = 0.019 (Figure 2 C). The HRV derived respiratory band power did not differ significantly between groups (Figure 3 C2, p = 0.916).
- **Cardiac band (0.5–1.5 Hz):** The fNIRS derived signal power remained unchanged between groups for all concentrations, indicating that cardiac-driven pulsations are preserved in OSA (Figure 2 D). HRV derived cardiac band showed no significant change between the groups, Figure 3 D2, p = 0.086).

#### HRV analysis

Analysis of Polar-derived HRV metrics from the same 30-minute windows used for fNIRS analysis (PWOSA_polar_ & HC_polar_) revealed a consistent increase in parallel in VLF_HRV_ power (Figure 3 E, Figure 4 A). PWOSA_Polar_ group exhibited significantly higher VLF_HRV_ power (0.003–0.04 Hz) compared to control subjects (p = 0.007). This finding parallels the elevated VLF_fNIRS_ power observed in fNIRS signals. Despite the significant differences in power with only at VLF_HRV_ range, the different groups have different characteristics across the spectrum (Figure 5) In contrast, other HRV frequency domain measures did not differ significantly between the groups: LF power (0.04–0.15 Hz, p = 0.434), HF power (0.15–0.4 Hz, p = 0.949), and total power (p = 0.112) (Figure 4 B,C,D). The LF/HF ratio, an index of sympathovagal balance, was also unchanged (p = 0.409).

**Figure 4.**
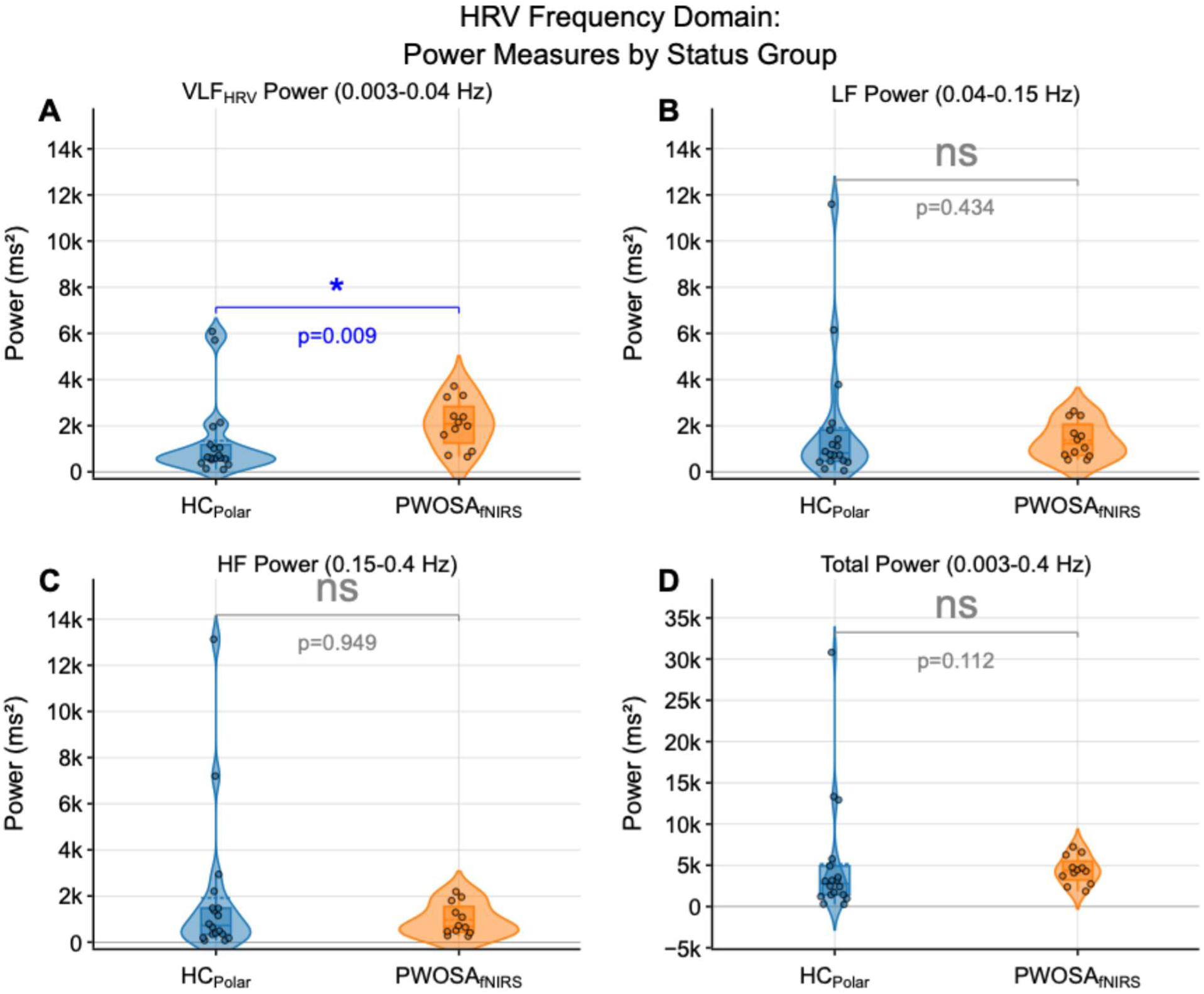
Heart rate variability frequency domain power in HC_polar_ and PWOSA_polar_ during 30-minute segments synchronized with fNIRS recordings. VLF_HRV_ power (0.003–0.04 Hz) was significantly elevated in OSA patients (p< 0.05), paralleling the fNIRS findings.

**Figure 5.**
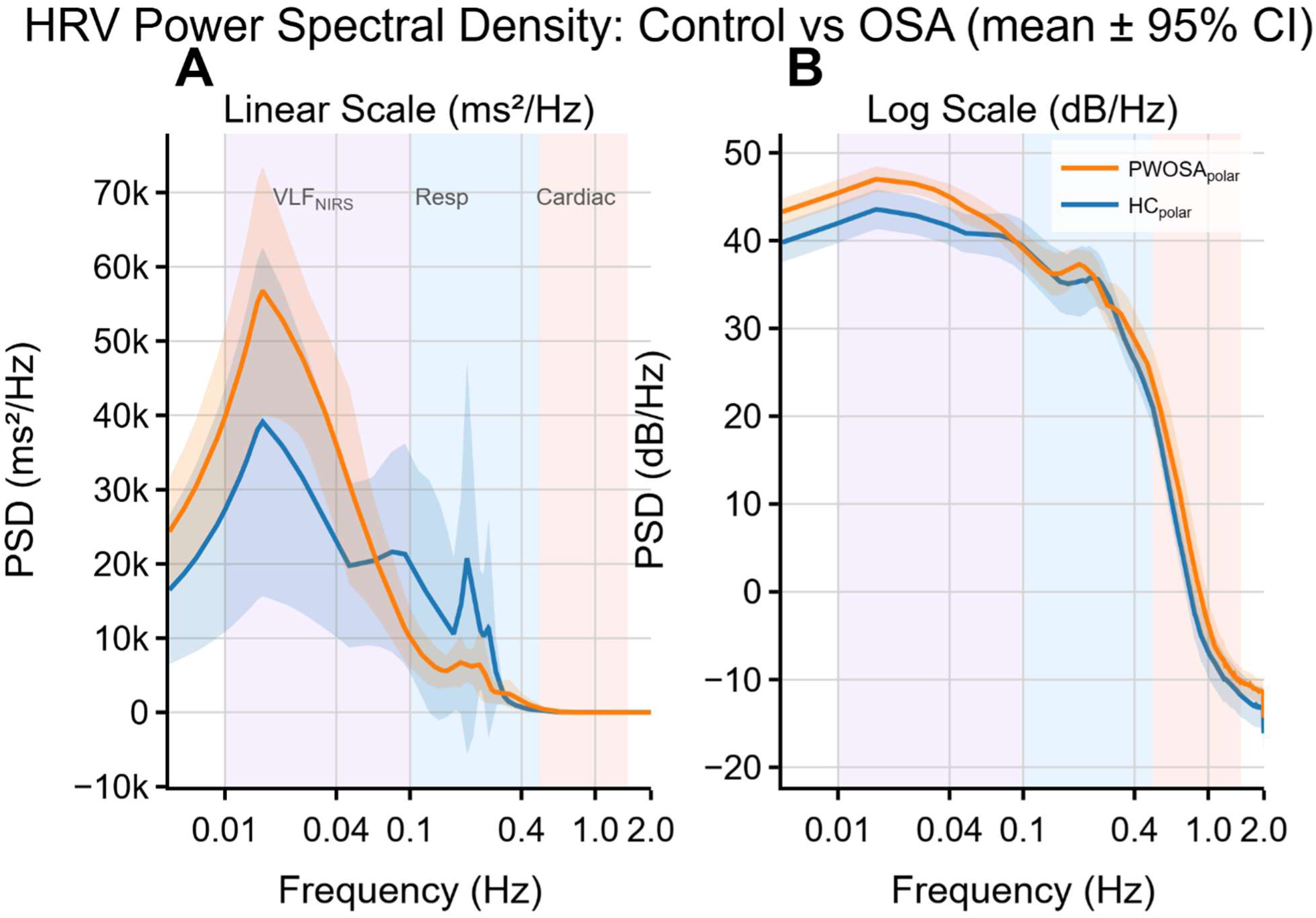
Power spectral densities between HC_polar_ and PWOSA_polar_ groups. A = linear scale, B = Logarithmic scale

**Time domain analysis** showed that RMSSD during the 30-minute segments did not differ significantly between groups (p = 0.842), in contrast to the event based whole-night analysis presented (Figure 10).

#### Hypoxic burden

Within PWOSA_fnirs_ group in 30min sleep segments, the hypoxic burden was significantly correlated with the power on cardiac band in CSF (r = 0.78, p = 0.003), HbT (r = 0.76, p = 0.004) and HbO (r = 0.75, p = 0.005) (Figure 6). No significant correlations were observed between hypoxic burden and the remaining concentration signal power bands (Figure 6). Time under 90% oxygen saturation was also positively and significantly in correlation of the power in cardiac band with CSF (r = 0.73, p = 0.007), HbR (r = 0.70, p = 0.011) and in VLF_fNIRS_ with HbR (r = 0.59, p = 0.042).

**Figure 6.**
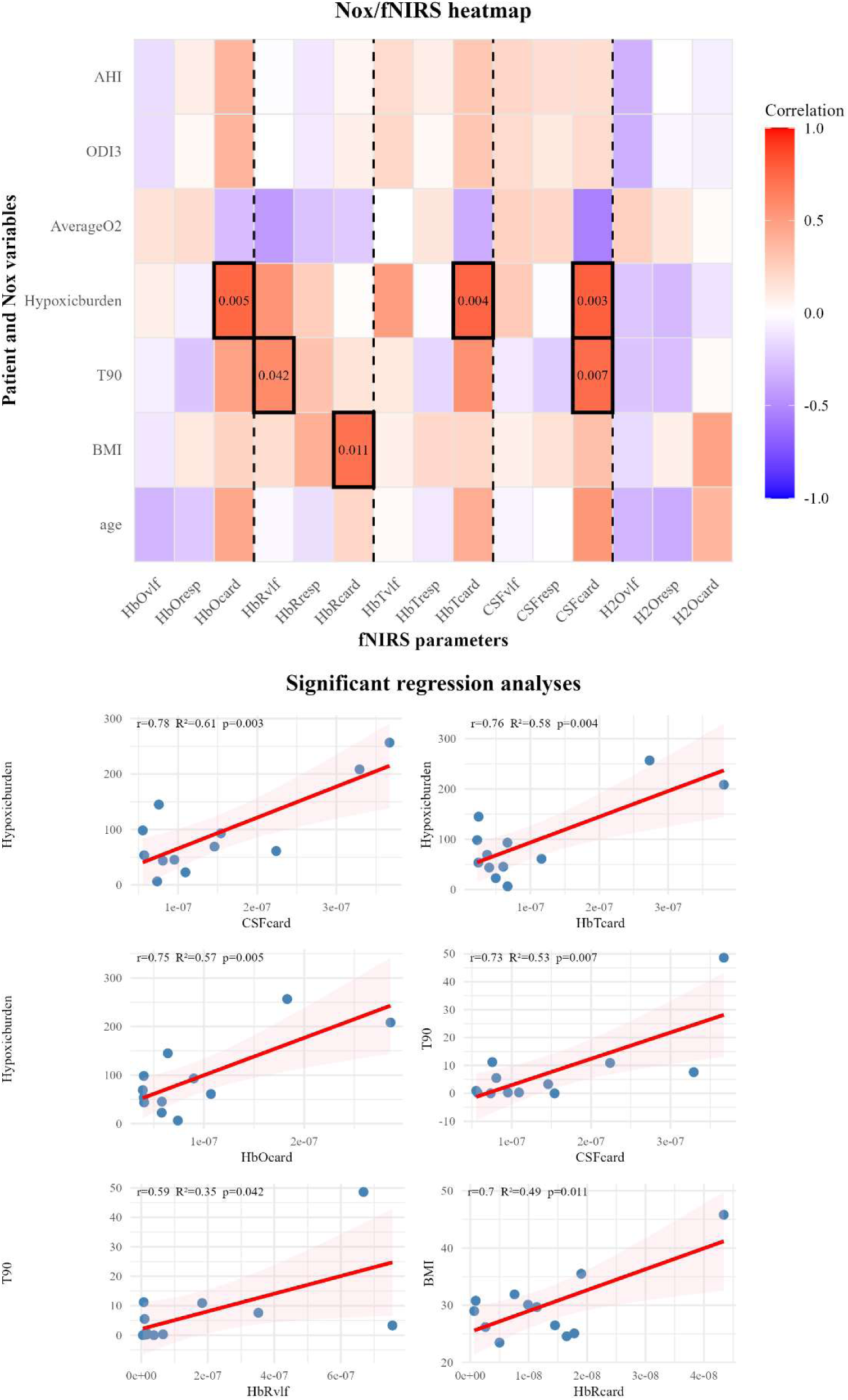
Correlations between sleep and clinical parameters and fNIRS derived spectral band powers in PWOSA_nirs_ during high-quality sleep segments. (Top) Heatmap of Pearson correlation coefficients (r) between subjects and Nox-derived variables and demographics (age, BMI, Time under 90% oxygen saturation (T90), hypoxic burden, average SpO_2_, Oxygen desaturation index (ODI) and AHI) and fNIRS spectral powers across five chromophore signals (HbO, HbR,HbT,CSF,H_2_O) and three frequency bands (VLF_fNIRS_, respiratory, cardiac). Significant correlations (p<0.05) are outlines in black and labelled with their p-values. (Bottom) Regression plots for all significant correlations; shaded regions show 95% confidence interval of the regression fit.

#### Coherence

Out of the 13 compared concentration signal pairs, five reached statistical significance. All the results are presented in supplementary Table 1. In this section we focus on the concentration signal pairs that reached significance. In PWOSA_fnirs_ group inter-signal synchrony was reduced compared to control group. This reduction was observed across multiple concentration pairs and frequency pairs (Figure 7). CSF/ HbO coherence was reduced in respiratory band (HC_fnirs_ = 0.898, PWOSA_fnirs_ = 0.776, p = 0.019, Figure 7 E) and also in cardiac band (HC_fnirs_ = 0.918, PWOSA_fnirs_ = 0.811, p = 0.037, Figure 3 C). HbT/HbO coherence was similarly reduced in respiratory band (HC_fnirs_ = 0.868, PWOSA_fnirs_ = 0.756, p = 0.028, Figure 7D) and also in cardiac band (HC_fnirs_ = 0.843, PWOSA_fnirs_ = 0.711, p = 0.025, Figure 7B).

**Figure 7.**
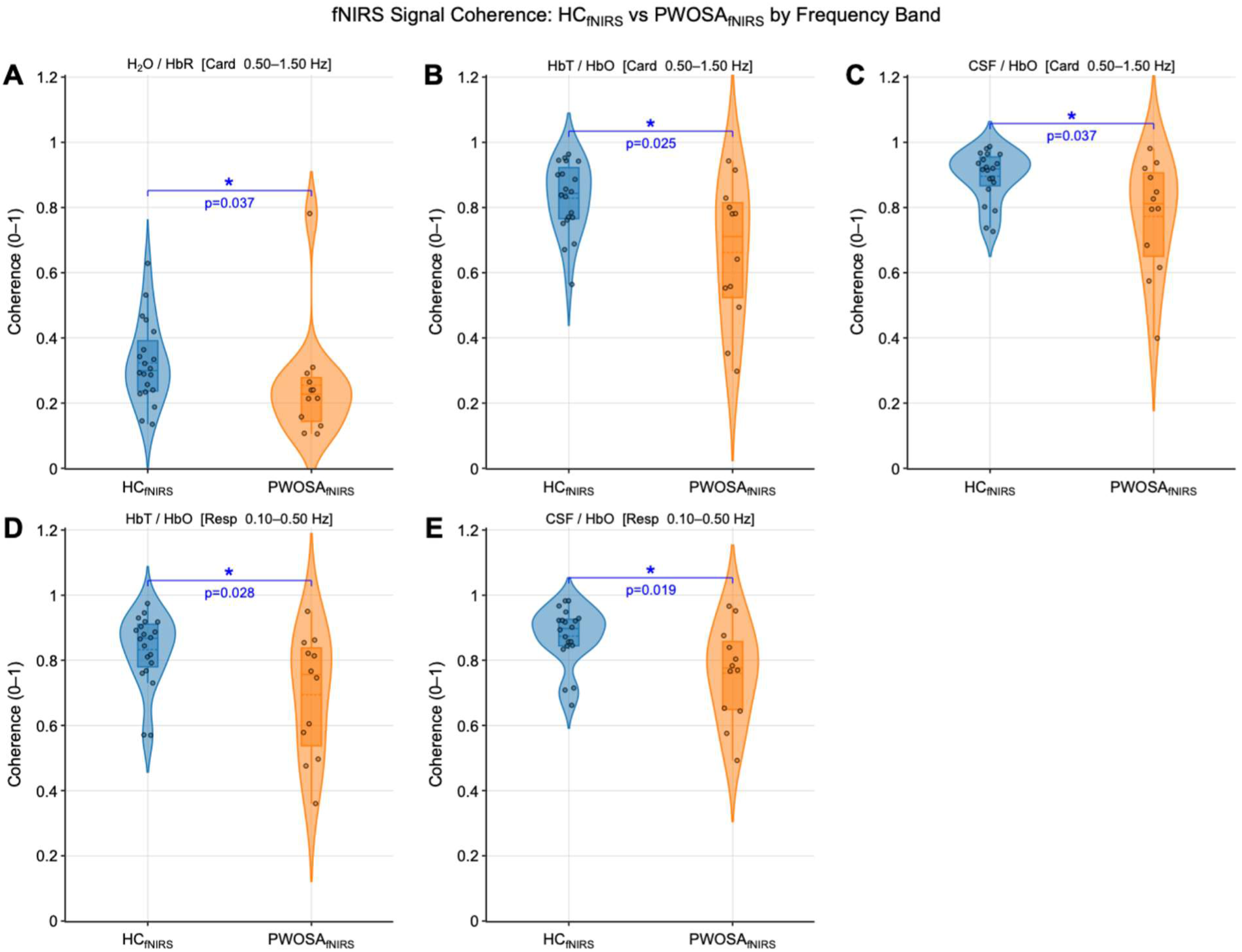
fNIRS inter-signal coherence in healthy controls (HC_fnirs_, blue) and OSA patients (PWOSA_fnirs_, orange). Magnitude-squared coherence was averaged within each frequency band and compared between groups using the Mann-Whitney U test. Asterisks denote p<0.05; ns =not significant. Panel A shows H2O/HbR in cardiac band, panels B&D show HbT/HbO at cardiac and respiratory bands and panels C&E shows CSF/HbO coherence across respiratory, and cardiac bands.

Also, the H_2_O/HbR coherence in the cardiac band was significantly lower in patients (HC_fnirs_ = 0.300, PWOSA_fnirs_ = 0.227, p = 0.037, Figure 7A). However, the absolute values were clearly lower than the other pairs. Together these findings demonstrate a broad reduction in the oscillatory synchrony between hemodynamic compartments during sleep in OSA.

#### Phase Transfer Entropy

Out of the 13 compared concentration pairs, only five showed significant differences in pTE between the groups. All the results are presented in supplementary Table 1. The pTE analysis revealed disrupted directionality of cardiac frequency coupling in OSA patients (Figure 8). In HC_fnirs_ net information flow was consistently directed toward the deoxygenated haemoglobin compartment across all four cardiac-band pairs (Figure 8A-D). In PWOSA_fnirs_ this directionality was absent or reversed. Net information flow toward HbR was significantly reduced from CSF (HC_fnirs_ = 0.089, PWOSA_fnirs_ = 0.009, p = 0.034, Figure 8A) and also from HbT (HC_fnirs_ = 0.096, PWOSA_fnirs_ = 0.006, p = 0.019, Figure 4 C). Furthermore, the directionality of coupling was reversed for HbO to HbR (HC_fnirs_ = 0.053, PWOSA_fnirs_ = -0.023, p = 0.023, Figure 8 B) and for H_2_O to HbR (HC_fnirs_ = 0.039, PWOSA_fnirs_ = -0.021, p = 0.012, Figure 8 D). Additionally, the information flow form CSF to HbO at respiratory band was significantly reduced (HC_fnirs_ = 0.0042, PWOSA_fnirs_ = - 0.0035, p = 0.045, Figure 8 E).

**Figure 8.**
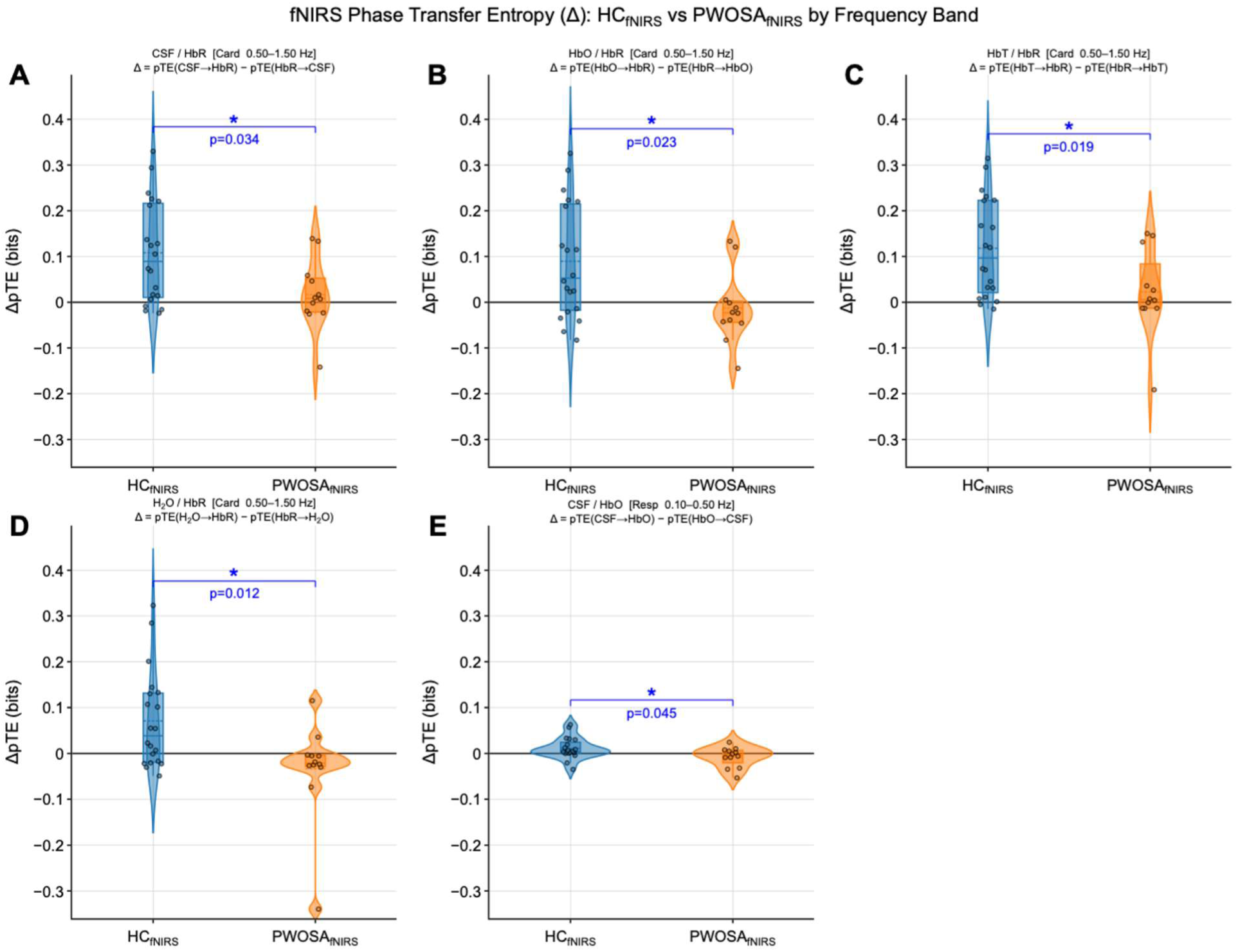
Net phase transfer entropy (ΔTE = pTE(X→Y) − pTE(Y→X)) between fNIRS concentration signal pairs in HC_fnirs_ (blue) and PWOSA_fnirs_ (orange). Positive ΔTE indicates net directed information flow from the first to the second signal. Panels A–D show cardiac-band pairs involving HbR; panels E–F show VLF_fNIRS_-band pairs involving HbT. Asterisks denote p<0.05.

### 3.3 Event-Based Whole-Night Analysis

The following results are derived from Polar recordings across the entire night for HC_polar_ & PWOSA_polar_ study groups.

#### Heart Rate During Sleep Events

Heart rate patterns differed significantly between OSA and control groups across all analyzed events. During event-free segments, the HC_polar_ group demonstrated a median heart rate of 63 bpm (interquartile range IQR 56–71 bpm) while PWOSA_polar_ group showed significantly higher median heart rate of 67 bpm (IQR 60–74 bpm; p *<* 0.001).

During obstructive respiratory events, PWOSA_polar_ group exhibited a median heart rate of 66 bpm (IQR 59–73 bpm) and during desaturation, the respective values were 66 bpm (IQR 60–71 bpm). All showed significant differences compared to the control group (p *<* 0.001; Figure 9).

**Figure 9.**
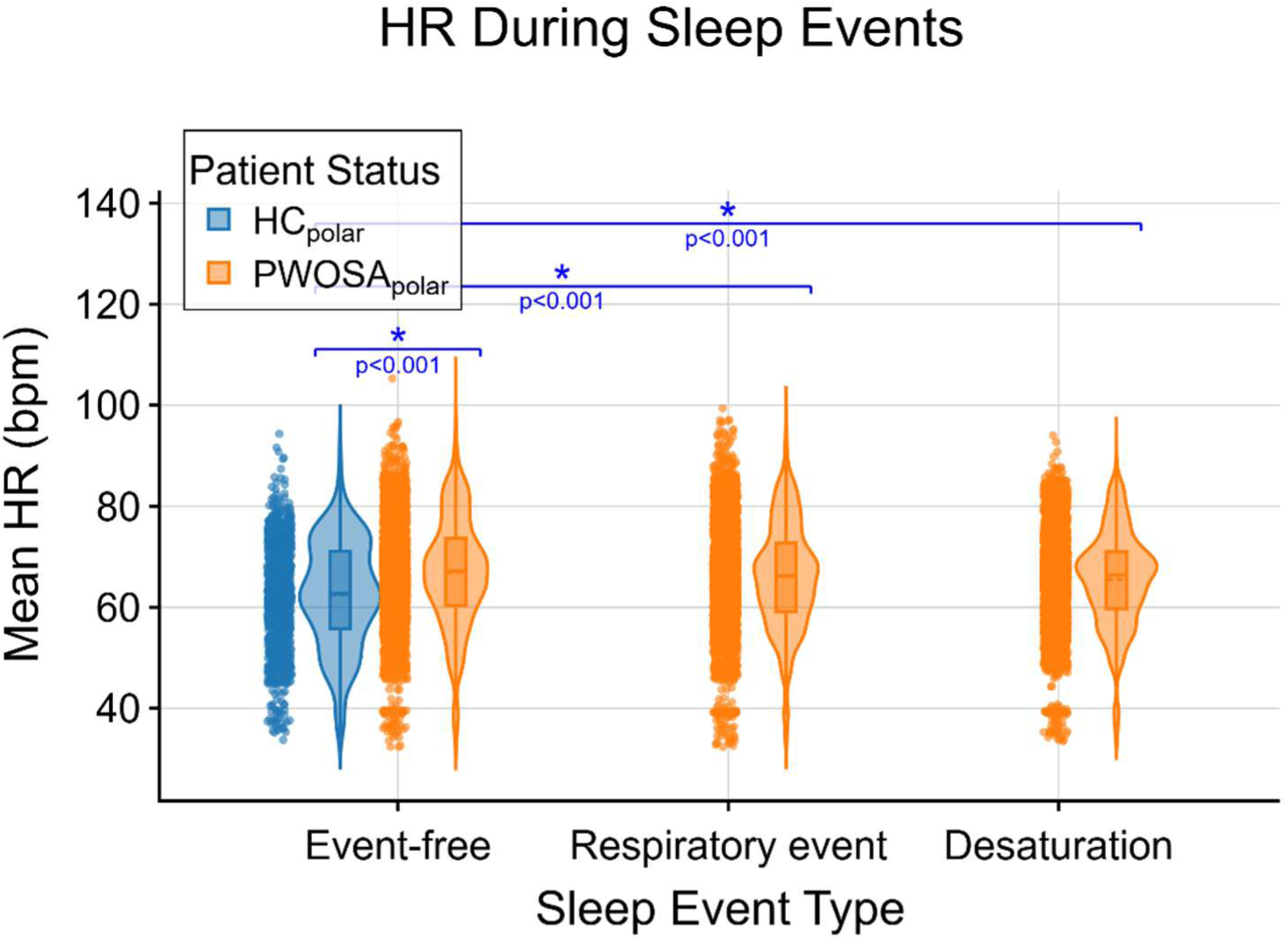
Heart rate during events in HC_polar_ and PWOSA_polar_ study groups. PWOSA_polar_ showed significantly elevated heart rate across all event types, including event-free periods, respiratory and desaturation events (all p < 0.001)

#### Heart Rate Variability (RMSSD)

HRV levels differed significantly between PWOSA_polar_ group and HC_polar_ group in all analyzed events (Figure 10). Even during event-free periods, the control group maintained median RMSSD of 43.0 ms (IQR 28.6–70.3 ms) while OSA patients showed significantly lower median RMSSD of 32.5 ms (IQR 23.4–45.6 ms; p *<* 0.001). The same pattern was observed during obstructive respiratory events, and desaturations (all p *<* 0.001).

**Figure 10.**
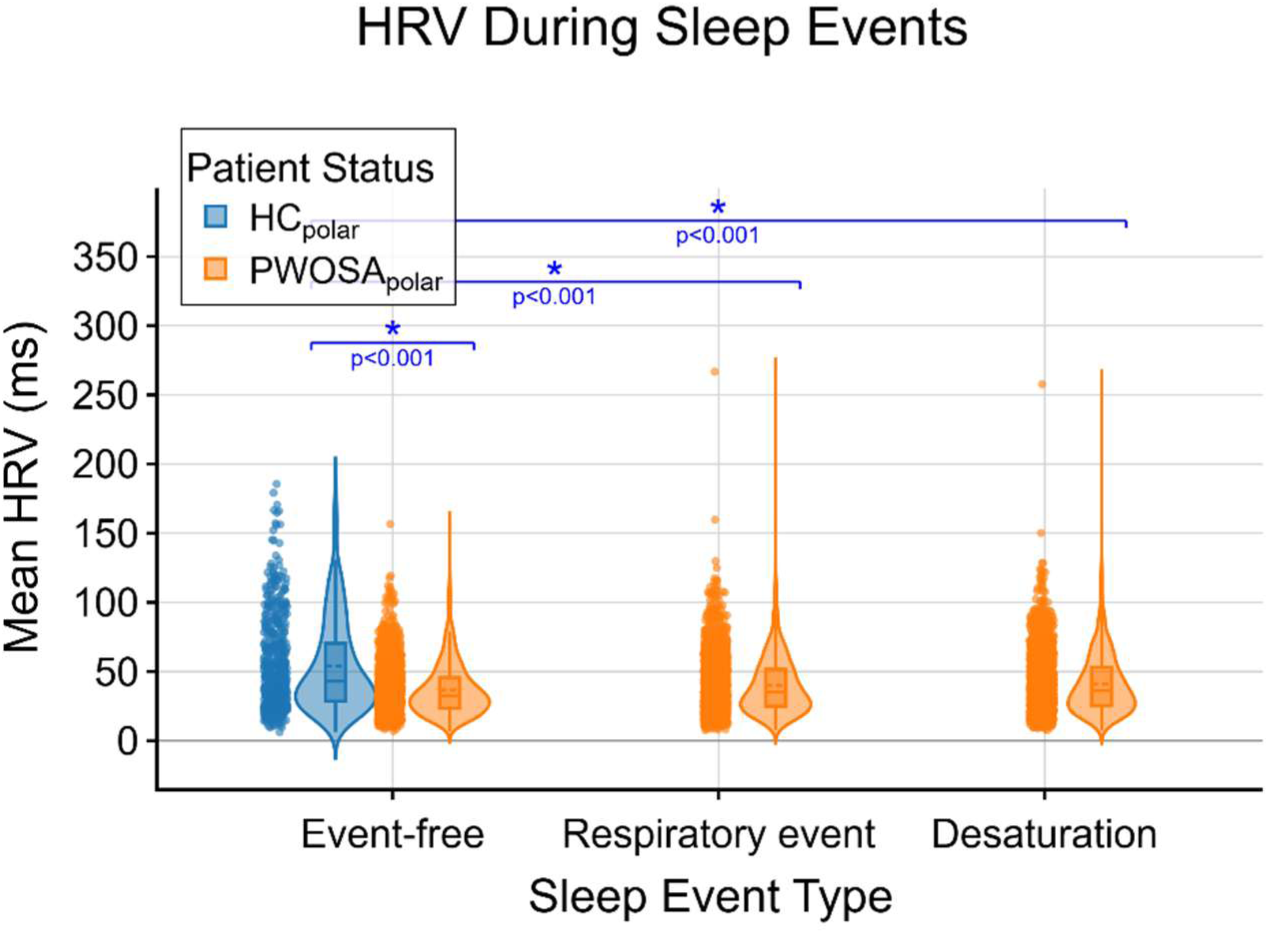
Heart rate variability (RMSSD) during events. PWOSA_polar_ exhibited significantly reduced RMSSD compared to HC_polar_ across all event types (all, p < 0.001), indicating decreased parasympathetic tone that persists beyond acute respiratory events

#### VRRI Analysis

During event-free periods, HC_polar_ group exhibited a median VRRI of 0.0 units (IQR -0.01 to 0.01) while PWOSA_polar_ group showed a median VRRI of 0.0 units (IQR -0.02 to 0.02). The difference was non-significant (p = 0.553), as expected given the nature of the VRRI parameter during non-event periods. The largest difference was observed during obstructive events, where PWOSA_polar_ group had a median VRRI of -0.04 units (IQR -0.07 to -0.007), with significant difference compared to event-free periods (p *<* 0.001). Hypopnea and desaturation events also showed significant differences (Figure 11).

**Figure 11.**
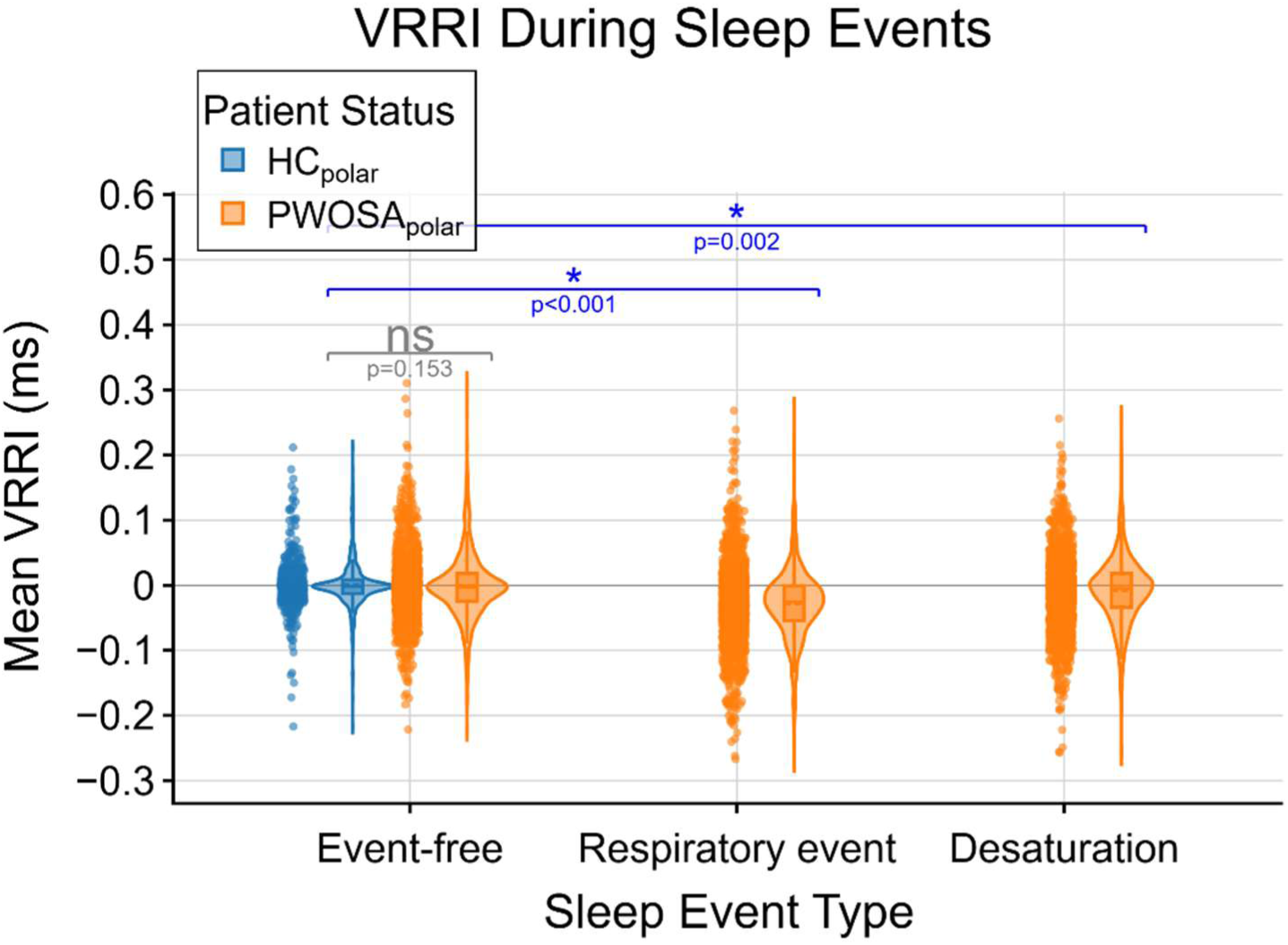
VRRI (depth of valley of RRI fluctuation) during events. During event-free periods, no significant difference was observed between groups (p = 0.153). However, during respiratory events (p<0.001), and desaturation events (p<0.05), PWOSA_polar_ group showed significantly altered VRRI values, reflecting acute autonomic responses to respiratory obstruction

#### Bioimpedance Phase Angle

Analysis of bioimpedance phase angle during overnight recordings showed differences between groups across multiple sleep event types. During event-free periods, the mean phase angle was 20.88^◦^ ± 10.70^◦^ for HC_polar_ group and 23.16^◦^ ± 8.26^◦^ for PWOSA_polar_ group (p =0.000). During respiratory events, the corresponding value was 22.61^◦^ ± 9.38^◦^, with significant difference compared to HC_polar_ groups event-free periods (p = 0.003). Desaturation events also showed significant differences from the HC_polar_ group with 23.12^◦^ ± 8.25^◦^ (p= 0.000) (Figure 12).

**Figure 12.**
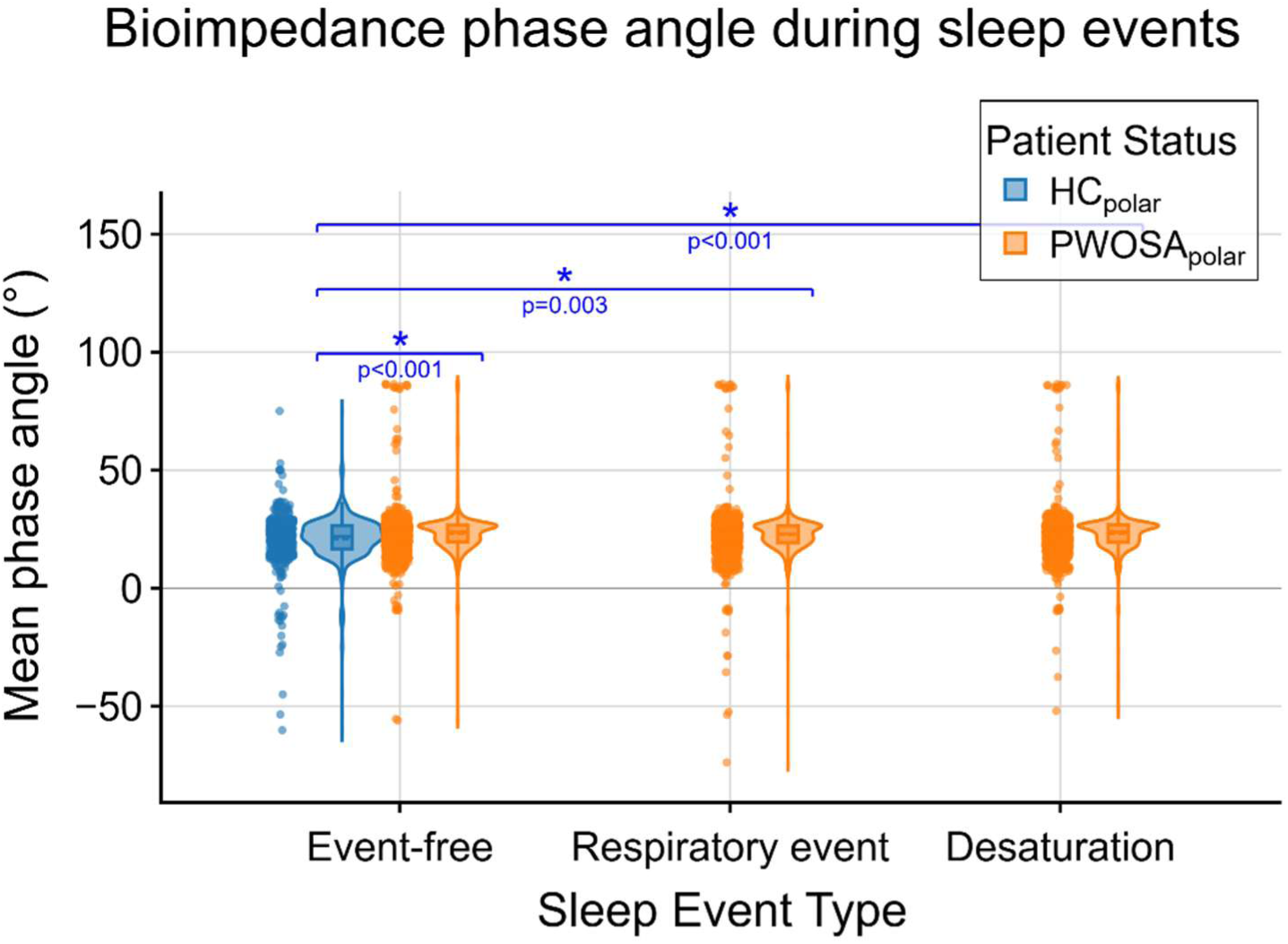
Bioimpedance phase angle during events. PWOSA_polar_ showed significantly different phase angles during respiratory events (p<0.05) and desaturation (p< 0.001) compared to HC_polar_ groups event-free periods.

## 4 Discussion

This study demonstrates that OSA is associated with a measurable alteration in cortical neurofluid dynamics during sleep. These dynamics were characterized by reduced spectral entropy and increased power of very-low-frequency (VLF, 0.01–0.1 Hz) and respiratory-band (0.1–0.5 Hz) oscillations in fNIRS-derived HbO, HbR, HbT, CSF, and H₂O signals. In contrast, cardiac-band (0.5–1.5 Hz) power was preserved. Parallel alterations in heart rate variability (HRV), including reduced RMSSD and elevated nocturnal heart rate, indicate persistent autonomic imbalance, i.e. sympathetic tone increase and parasympathetic decrease, extending beyond discrete apneic events. Additionally, the differences between the study groups in phase entropy and coherence analysis show that OSA not only affects overall spectral properties but also relationships between different fNIRS derivatives. Together, these findings suggest that OSA is associated with a redistribution and simplification of physiological oscillatory dynamics during sleep affecting normal brain physiology.

During normal breathing cycle intrathoracic pressure gradient is usually oscillates between -5mmHg and +5mmHg (Neupane and Jamil, 2026), and usually during sleep these pressures are maintained on a similar scale. However, during sleep apnea, negative pressure could be lowered as low as -80mmHg (Kasai, 2012), causing, for instance increased left ventricular afterload and reduced cardiac output (Kasai, 2012), affecting overall physiology and causing well-known comorbidities (Lv et al., 2023). As obstructive apneas are characterized by repeated negative intrathoracic (Kasai, 2012) pressure swings during inhalation against a collapsed airway, which probably augment venous return fluctuations from the brain, alter cerebral venous pressure, and disturb oscillatory forces. Such mechanical negative pressure amplification may represent reduced inward pulsations of respiratory-driven hemodynamic oscillations, while increasing outward pulsations, causing vacuum-like motion to the brain, and thus disturbing usual glymphatic function. Thus, the prevalent pressure condition is totally different in normal breathing (between -5mmHg and +5mmHg) and OSA (down to -80mmHg), and breathing against pressure, e.g. Valsalva maneuver (increasing pressure up to +40mmHg (Yu et al., 2025)). However, because the spectral analyses does not account for temporal changes, the spectral power is similar for pressure gradients between -35mmHg and +5mmHg or between -20mmHg and +20mmHg. Therefore, the increased respiratory power does not necessarily imply improved fluid transport efficiency but rather, this excessive oscillatory forcing may instead reflect instability within the cerebrovascular system.

The increase in respiratory-band power likely reflects transmission of intrathoracic pressure oscillations to both the cerebral vasculature and the incompressible CSF within the spinal canal. During obstructive events, the increased inspiratory effort against negative intrathoracic pressure appears to alter brain hydrodynamics. Because these breathing interruptions occur periodically, glymphatic physiology may also be disrupted in a cyclical manner, thereby disturbing normal brain hydrodynamics and placing additional strain on sleep physiology. As respiratory brain pulsations influence CSF pulsatility, they may also affect brain waste clearance. In addition, OSA is well known to disrupt sleep, whereas sleep itself enhances glymphatic function, further supporting the potential physiological significance of these disturbances (Helakari et al. 2022; Xie et al. 2013; Roy et al. 2022; Sangalli and Boggero 2023; Fultz et al. 2019). Furthermore, these alterations may impair cerebral autoregulatory mechanisms in a manner analogous to peripheral vascular disturbances (Olopade et al. 2007; Ilvesmäki et al. 2024).

Similarly, the increase in VLF_fNIRS_ power suggests altered slow vasomotor regulation. VLF_fNIRS_ oscillations in cerebral hemodynamics are commonly linked to endothelial and neurogenic modulation of vascular tone and are influenced by sympathetic activity (Andersen et al., 2018; Helakari et al., 2022; Kiviniemi et al., 2016). In OSA, repetitive hypoxia and arousal-related sympathetic activation may cause higher-amplitude vasomotor oscillation. Whether this reflects compensatory autoregulatory activation or reduced regulatory precision cannot be conclusively determined from spectral amplitudes alone. However, this increased VLF _NIRS_ power may indicate a failure of the cerebral windkessel mechanism to damp slow frequency pressure swings, allowing excessive pulsatile energy to be transmitted into microcirculation. The reduction in entropy also supports the interpretation that oscillatory activity becomes more stereotyped and less dynamically distributed across frequencies.

In contrast, cardiac-band power remained unchanged between groups, although it is common knowledge that OSA patients pulse increase during respiratory event (Arikawa et al., 2020). Cardiac pulsatility is a fundamental driver of brain pulsations and perivascular fluid displacement (Iliff et al., 2012; Kiviniemi et al., 2016), yet its spectral amplitude may be relatively preserved even in the presence of pathological vascular modulation. In normal sleep card pulsation drops compared to awake state probably minimizing cardiac pulsations effect during sleep (Helakari et al., 2022; Kiviniemi et al., 2017; Tuunanen et al., 2024). Although, it is probable that also reduced cardiac output affects these pulsations and might even compensate each other. Hence, preservation of cardiac-band power does not imply intact coupling efficiency or effective propagation of pulsatile energy through parenchyma. The correlation between hypoxic burden and cardiac band HbO and HbT power may reflect the autonomic nervous system activation that is associated with cumulative nocturnal hypoxia (Azarbarzin et al., 2020). The corresponding correlation with CSF could however suggest that intracranial changes in OSA patients may be sensitive to overall hypoxic load rather than discrete apnea events alone (Coso et al., 2024).

Coherence analysis revealed reduced synchronization between vascular and fluid-related oscillations across cardiac and respiratory bands in OSA, particularly in the CSF/HbO and HbT/HbO pair selected to probe the CWK effect (Oliveira et al., 2025; Wagshul et al., 2011; Westerhof et al., 2009). Concurrently, phase transfer entropy analysis showed that the normal directional coupling toward HbR at cardiac frequencies was disrupted in OSA, and the CSF/HbO coupling in the respiratory band was reduced. Together, these findings suggest a loss of coordinated oscillatory organization rather than a simple increase of pulsatile drivers disturbing the usual glymphatic function. Such desynchronization may reflect impaired and disease-related vascular compliance, altered autoregulatory function, or disrupted transmission of pulsatile energy between compartments. These disruptions are hallmark features of CWK dysfunction, where a reduction in craniospinal compliance leads to an impaired ability to buffer intracranial pressure waves across multiple frequency bands. These findings also suggest that OSA selectively disrupts directed coupling mechanisms operating at cardiac frequencies within the deoxygenated hemodynamic compartment.

The autonomic data provides convergent indication of systemic autonomic dysregulation. Reduced RMSSD and persistently elevated heart rate during whole night sleep, demonstrate sustained sympathetic predominance and diminished parasympathetic modulation (Task Force ESC/NASPE, 1996). Chronic sympathetic activation is known to influence vascular tone, arterial stiffness, and endothelial function, potentially contributing to the altered VLF_HRV_ dynamics observed in cortical signals (Andersen et al., 2018). The parallel increase in VLF_HRV_ power within HRV metrics further supports the presence of systemic slow-rhythm dysregulation. However, VLF_HRV_ oscillations in HRV and cerebral hemodynamics arise from partially distinct physiological sources. Thus, currently these findings should be interpreted as parallel manifestations of autonomic imbalance rather than direct evidence of shared oscillators. VRRI analysis showed a negative valley PPI valley in PWOSA_polar_ during respiratory events (median -0.04 vs 0.0 in HC_polar_, p<0.001) reflecting the acute parasympathetic withdrawal-rebound cycle at apnea offset (Arikawa et al., 2020). While average HRV metrics capture the sustained autonomic imbalance, VRRI captures the brief reactions tied to individual respiratory event.

As an exploratory marker, PhA differed significantly between groups during event-free, respiratory and desaturation periods. Because PhA is sensitive to cellular membrane integrity and tissue hydration but also to age and body composition (Barbosa-Silva et al., 2005; Holder, 2004), the baseline difference cannot be attributed to OSA alone. The event-related modulation is nonetheless consistent with acute physiological disturbations of OSA influencing tissue properties (Lv et al., 2023). As an exploratory finding, this result should be treated as preliminary and followed up in studies where OSA patients and controls are better matched in age and BMI.

Given that glymphatic transport has been proposed to depend on coordinated cardiac, respiratory, and vasomotor pulsations, disruption of multi-frequency balance may have implications for sleep-dependent fluid exchange (Fultz et al., 2019). Experimental models suggest that perivascular transport relies not solely on the magnitude of individual drivers but on their coordinated interaction (Fultz et al., 2019) and the compliance properties of the surrounding tissue (Kedarasetti et al., 2020). The observed redistribution of oscillatory power and reduction in complexity together with altered coordination between pulsations, are consistent with a state in which multi-frequency coupling becomes less efficient. Nevertheless, direct measures of CSF flux or tracer clearance were not obtained; therefore, impaired glymphatic function remains a physiologically plausible but unconfirmed interpretation.

### Technical considerations, strengths, limitations and future considerations

It should be noted that presented fNIRS methods measures cerebral hemoglobin concentration changes, total H_2_O including water bound with blood, and CSF dynamics mainly associated to the subarachnoid space (Myllylä et al 2018), hence, it does not directly quantify CSF flow or glymphatic function. Thus, the observed findings reflects decreased temporal irregularity and physiological multi-frequential complexity in cortical brain hydrodynamics rather than a direct measure of impaired clearance. In biological systems, reduced entropy is usually interpreted as diminished adaptive capacity, reflecting constrained regulatory dynamics (Lipsitz and Goldberger, 1992). Within this framework, the findings suggest that OSA is associated with a loss of flexible vascular modulation across interacting physiological frequencies. The loss of complexity, combined with the observed power redistribution, points toward a rigid system where the CWK mechanism’s intrinsic protective damping is exhausted, diminishing the brain’s adaptive capacity.

Further limitations warrant consideration. First, fNIRS measurements are restricted to cortical regions and cannot assess deep structures such as the hippocampus, which are highly relevant to OSA-related cognitive decline (Ferrari and Quaresima, 2012; Torelli et al., 2011). Second, the optical measurement with fNIRS provides only indirect estimates of cerebral hemodynamic and water dynamics; it does not directly quantify intracranial compliance or CSF displacement (Myllylä et al., 2018). Third, the current study design precludes inference regarding causality or reversibility with OSA treatment. Additionally, because the analyzed periods were selected primarily based on data quality criteria and sleep staging was not available, differences in sleep stages between subjects cannot be interpreted. However, this limitation applies similarly to both groups. Although the exact sleep stages remain unknown, the measurement setup clearly indicates that the subjects were asleep during the analyzed data segments. Finally, potential confounders including body mass index, blood pressure, and medication use may influence vascular oscillatory behavior.

It is also notable that RMSSD differed between groups in the event-based whole-night analysis (32.5 ms vs 43.0 ms, p < 0.001), but not in the matched 30-minute high-quality segments (p = 0.842), despite the high density of respiratory events with PWOSA group (segment AHI 44.7 ± 23.7). This dissociation likely reflects how each analysis samples the night: averaging RMSSD over a 30-minute window mixes event and event-free periods, masking the brief autonomic changes that differ between groups, whereas event-based analysis focuses on the periods where these changes occur.

Additionally, it is important to consider, that reduced entropy could also arise from spectral dominance of frequency bands – such as exaggerated respiratory oscillations – rather than global simplification of underlying dynamics. Furthermore, differences in sleep stage distribution may further influence oscillatory properties, as slow-wave sleep is known to modulate vascular and interstitial dynamics (Xie et al., 2013). While our analysis controlled for major confounders, residual effects cannot be excluded.

Future studies integrating multimodal imaging, event and sleep stage related analyses, or longitudinal assessment before and after different treatment strategies will be essential to determine whether normalization of physiological pulsations complexity parallels improvements in measurable variables. This would clarify whether altered pulsatility represents an epiphenomenon of autonomic stress or a contributory factor in sleep-related vulnerability.

#### 4.1 Conclusions

The present findings provide clear evidence that OSA is associated with a reorganization of physiological brain pulsations during sleep. Rather than a uniform increase in pulsatile activity, OSA appears to produce a selective redistribution of oscillatory energy across frequencies, accompanied by chronic autonomic imbalance. These alterations may represent a mechanistic link between sleep fragmentation, cerebrovascular dysregulation, and the increased risk of neurodegenerative pathology observed in OSA populations.

Furthermore, this study demonstrates that fNIRS based spectral analysis can non-invasively detect these cerebral alterations. When combined with wearable HRV monitoring, this approach offers a promising, scalable bedside tool for assessing the impact of sleep disorders on brain health and monitoring the physiological efficacy of treatments.

## 4.2 Acknowledgements

We wish to thank all study subjects whose attendance made this study possible. We also thank Hany Ferdinando, Martti Ilvesmäki and Jari Paunonen for fNIRS related introduction and necessary first level preprocessing steps. Finally, we offer our thanks to the funders of this study: Research Council of Finland (360508) (JK), Finnish Medical Foundation (J.K.), Pohjois-Suomen terveydenhuollon tukisäätiö, Terttu-Foundation (JK), BF (11,146/31/2022) (TM) and State Research Funding, VTR (JK). We also thank Polar Electro Oy for providing wearable devices for the study.

## 4.3 Declaration of generative AI use

Artificial intelligence tools (Gemini 3, Claude Opus 4.5, Claude Sonnet 4) were used to assist with data analysis pipelines. However, all final analyses, interpretation of results, and scientific conclusions were composed, verified, and approved solely by the authors.

## 4.4 Data availability statement

The individual data from this study cannot be shared because of privacy issues of clinical data. Data available upon request from the corresponding author for research only after ethical approval for the specific study.

---

**Supplementary Table 1.** coherence and pTE results. Card, cardiac band (0.50–1.50 Hz); Resp, respiratory band (0.10–0.50 Hz); VLF, very low frequency band (0.01–0.10 Hz). Direction shows the dominant transfer direction: Δ = pTE(X→Y) − pTE(Y→X). * p < 0.05.

| Measure | Pair | Band | Direction | Ctrl med. | Pt med. | p | Sig. |
| --- | --- | --- | --- | --- | --- | --- | --- |
| Coherence | CSF / HbR | Card |  | 0,2121 | 0,1559 | 0,2058 |  |
| Coherence | HbO / HbR | Card |  | 0,3557 | 0,2931 | 0,4027 |  |
| Coherence | HbT / HbR | Card |  | 0,1800 | 0,1606 | 0,5463 |  |
| Coherence | Water / HbR | Card |  | 0,2996 | 0,2272 | **0,0373** | * |
| Coherence | HbT / HbO | Card |  | 0,8434 | 0,7109 | **0,0252** | * |
| Coherence | CSF / HbO | Card |  | 0,9182 | 0,8113 | **0,0373** | * |
| Coherence | HbT / CSF | Resp |  | 0,9070 | 0,8978 | 0,5207 |  |
| Coherence | HbT / Water | Resp |  | 0,5730 | 0,6286 | 0,5992 |  |
| Coherence | HbT / HbO | Resp |  | 0,8679 | 0,7562 | **0,0279** | * |
| Coherence | CSF / HbO | Resp |  | 0,8978 | 0,7763 | **0,0185** | * |
| Coherence | HbT / CSF | VLF |  | 0,9838 | 0,9509 | 0,7407 |  |
| Coherence | HbT / Water | VLF |  | 0,8406 | 0,7685 | 0,8610 |  |
| Coherence | HbO / HbR | VLF |  | 0,2370 | 0,1821 | 0,3811 |  |
| $\Delta pTE$ | CSF / HbR | Card | CSF → HbR | 0,089336 | 0,008529 | **0,0339** | * |
| $\Delta pTE$ | HbO / HbR | Card | HbO → HbR | 0,052640 | -0,022983 | **0,0228** | * |
| $\Delta pTE$ | HbT / HbR | Card | HbT → HbR | 0,096496 | 0,005897 | **0,0185** | * |
| $\Delta pTE$ | Water / HbR | Card | Water → HbR | 0,038598 | -0,021028 | **0,0121** | * |
| $\Delta pTE$ | HbT / HbO | Card | HbT → HbO | 0,027550 | 0,029229 | 0,7407 | |
| $\Delta pTE$ | CSF / HbO | Card | CSF → HbO | 0,013029 | 0,018899 | 0,4249 | |
| $\Delta pTE$ | HbT / CSF | Resp | HbT → CSF | 0,001698 | -0,000503 | 0,4958 | |
| $\Delta pTE$ | HbT / Water | Resp | HbT → Water | -0,001762 | -0,006828 | 0,5463 | |
| $\Delta pTE$ | HbT / HbO | Resp | HbT → HbO | 0,003710 | -0,000414 | 0,4958 | |
| $\Delta pTE$ | CSF / HbO | Resp | CSF → HbO | 0,004186 | -0,003526 | **0,0450** | * |
| $\Delta pTE$ | HbT / CSF | VLF | HbT → CSF | -0,001279 | 0,005918 | 0,2058 | |
| $\Delta pTE$ | HbT / Water | VLF | HbT → Water | -0,009872 | -0,008075 | 0,7407 | |
| $\Delta pTE$ | HbO / HbR | VLF | HbO → HbR | -0,041955 | -0,010172 | 0,9534 | |

